# Differential Hsp90-dependent gene expression is strain-specific and common among yeast strains

**DOI:** 10.1101/2020.11.19.389437

**Authors:** Po-Hsiang Hung, Chia-Wei Liao, Fu-Hsuan Ko, Huai-Kuang Tsai, Jun-Yi Leu

## Abstract

Enhanced phenotypic diversity increases the likelihood of a population surviving catastrophic conditions. It has been suggested that Hsp90, an essential molecular chaperone in eukaryotes, can suppress (i.e., buffer) or enhance (i.e., potentiate) the effects of genetic variation, enabling organisms to adjust their levels of phenotypic diversity in response to environmental cues. Many Hsp90-interacting proteins are involved in signaling transduction pathways and transcriptional regulation. However, it remains unclear if Hsp90-dependent differential gene expression is common in natural populations. By examining the gene expression profiles of five diverse yeast strains, we identified many genes exhibiting Hsp90-dependent strain-specific differential expression. We employed an analysis pipeline to identify transcription factors (TFs) potentially contributing to variable expression. We found that upon Hsp90 inhibition or heat stress, activities or abundances of Hsp90-dependent TFs varied among strains, resulting in differential strain-specific expression of their target genes, which consequently led to phenotypic diversity. We provide evidence that individual populations can readily display specific Hsp90-dependent gene expression, suggesting that the evolutionary impacts of Hsp90 are widespread in nature.

**Highlights:**

1. Hsp90-dependent gene expression varies among different yeast strains.
2. Hsp90 differentially influences transcriptional activity or protein abundances of transcription factors among yeast strains.
3. Differential strain-specific gene expression is correlated with phenotypic variations upon Hsp90 inhibition.
4. Hsp90-mediated strain-specific regulation manifests under environmental stress.

## Introduction

Living organisms constantly face the challenges of environmental change. Although most organisms are equipped with sophisticated regulatory mechanisms to acclimate to such change, devastating situations occasionally arise that can wipe out most individuals of a given population, leaving only a few individuals with diverse phenotypes to survive. Darwin’s evolutionary theory states that heritable phenotypic variation provides the foundation on which natural selection operates, allowing organisms to evolve (Darwin 1859). Nonetheless, maintaining phenotypic diversity may have fitness costs under typical growth conditions, especially if divergent phenotypes deviate from optimal fitness (Wright 1932). It is crucial for a population to maintain high fitness so that it can compete with other populations under normal conditions, but it must also harbor enough phenotypic diversity to allow it to survive under stressful conditions.

A balance between the requirements of population fitness and phenotypic diversity can probably be achieved through cryptic genetic variation, i.e., allowing variations that do not cause phenotypic change under normal conditions, but that can do so when environments or genetic backgrounds are altered (Rutherford 2000; Sangster et al. 2004). Due to its neutral phenotypic effect in normal conditions, a significant proportion of cryptic genetic variation accumulates in the populations probably through genetic drift, similar to other types of neutral mutations. However, under certain environments or genetic backgrounds in which the buffering effect is compromised, some cryptic variants can also be quickly selected and enriched if they provide beneficial effects.

Multiple mechanisms have been proposed to explain the on/off phenotypic effect of cryptic genetic variation (Jarosz et al. 2010; Lehner 2011). One of the mechanisms is the network hubs that work as capacitors to buffer phenotypic variation. Computational modeling has shown that complex gene networks can buffer the effects of genetic and environmental perturbations (Stearns et al. 1995; Bergman and Siegal 2003). Deleting the hub genes often leads to phenotypic diversity even in an isogenic yeast population, confirming the hub genes’ capacitor function (Levy and Siegal 2008). Among those network hubs, heat shock protein 90 (Hsp90) represents the best-characterized capacitor, which also raises several interesting and controversial issues in evolutionary and molecular biology (Siegal and Leu 2014).

The primary function of Hsp90 is to assist in protein folding or to maintain protein stability (Nathan et al. 1997). Therefore, it is suggested that Hsp90 may have the ability to partially or completely alleviate the effects of some mutations that affect protein conformation or stability when they occur in Hsp90 substrates (also called Hsp90 clients) (Sangster et al. 2004). These buffered mutations will have a chance to be maintained in the population as “cryptic” genetic variants. However, when Hsp90 is compromised by chemicals or recruited to perform other tasks under stress conditions, the phenotypic effect of these cryptic variants will be revealed. Many Hsp90 clients are involved in signaling transduction pathways and transcriptional regulation (Leach et al. 2012), further raising the possibility that the Hsp90 buffering (or capacitor) effect can have a strong influence on cell physiology and organism development.

The buffering effect of Hsp90 has been observed in a wide variety of organisms including yeast, plants, flies, fish, mice, and humans (Rutherford and Lindquist 1998; Sangster et al. 2008; Jarosz and Lindquist 2010; Hummel et al. 2017; Karras et al. 2017), suggesting that it is a conserved mechanism. However, most of the respective studies have focused solely on a single pathway or phenotype. In addition, a recent study in yeast cell morphology indicated that the effects of newly occurred mutations were more likely to be enhanced by Hsp90 (Geiler-Samerotte et al. 2016), suggesting the mutations that can be buffered by Hsp90 may not be as common as what we previously assumed. Thus, it had remained unclear how prevalent Hsp90-buffered targets are in genomes and if specific variants buffered by Hsp90 already exist in different populations. Without addressing these two issues, it is difficult to assess the real impacts of Hsp90-dependent buffering on evolution.

Other than its buffering effect, Hsp90 can also function as a potentiator to enhance rather than suppress mutation effects, i.e., phenotypic diversity is revealed under normal conditions but is reduced when Hsp90 is compromised (Cowen and Lindquist 2005). This potentiating effect of Hsp90 has been implicated in cancer development (Blagosklonny et al. 1995; An et al. 2000; Minami et al. 2002). However, just as for Hsp90-dependent buffering, little is known about Hsp90-dependent potentiation in natural populations, apart from the results of two yeast studies (Jarosz and Lindquist 2010; Geiler-Samerotte et al. 2016).

The budding yeast, *Saccharomyces cerevisiae*, provides an ideal system for studying Hsp90-dependent buffering and potentiating events at the population level. In the yeast proteome, Hsp90 physically interacts with more than 500 proteins (Oughtred et al. 2016). Whole genomes of divergent yeast strains isolated from various ecological environments and geographic locations have been sequenced, and their population structures are well understood (Liti et al. 2009; Skelly et al. 2013; Bergstrom et al. 2014). Moreover, standing genetic variation may have previously experienced natural selection, which would purge out the ones with deleterious phenotypic effects (Geiler-Samerotte et al. 2016). Therefore, the buffered genetic variation observed in populations will provide additionanl information that cannot be extracted from newly occurred mutations in laboratory evolution experiments.

Since many potential Hsp90 clients are transcription factors (TFs) (Taipale et al. 2012), we assumed that the buffering and potentiating effects of Hsp90 could be revealed by differences in the transcriptional profiles of multiple strains. We examined transcriptomes in response to compromised Hsp90 among five highly diverged yeast strains. We identified several hundred Hsp90-dependent genes that were differentially expressed in each strain. Using comparative genomics and gene expression profiling, we identified the candidate TFs associated with strain-specific gene expression variation. We experimentally demonstrate that differential activities or abundances of those TFs displaying strain-specific Hsp90-dependent regulation could further lead to phenotypic variation among strains.

## Results

### Altered Hsp90-dependent gene expression is prevalent among yeast strains

Many known clients of Hsp90 are TFs or kinases involved in signal transduction (Taipale et al. 2012). If Hsp90-buffered or -potentiated genetic variation occurs in those clients, it could result in differential gene expression of their downstream targets when Hsp90 activity is compromised. To identify population-level events buffered or potentiated by Hsp90, we specifically focused on strain-specific differential gene expression. We chose five yeast strains—laboratory strain S288C (LAB), a strain collected from a Japanese sake distillery (SK), as well as strains from Malaysia (ML), North America (NA), and West Africa (WA) (Liti et al. 2009)—to examine their transcriptomes in the absence or presence of the Hsp90 inhibitor Geldanamycin (GdA). These strains differ from each other by about 50,000-85,000 single nucleotide polymorphisms (SNPs), indicating that they are highly diverged. Among them, the NA and SK strains show the smallest pairwise genetic distance, while the LAB and ML strains show the longest distance (see Methods and Supplemental Fig. S1A). However, they all expressed similar levels of Hsp90 under normal growth conditions (Supplemental Fig. S2).

When we analyzed their transcriptomes by RNA-seq, we discovered that expression levels of hundreds to more than one thousand genes were altered upon GdA treatment (fold change > 2 or < 0.5, adjusted p-value < 0.05, Supplemental Fig. S3 and Table S1). Interestingly, only a small number of the differentially expressed genes (n=83) were shared by all five strains (Supplemental Fig. S4). To ensure that the variation in expression profiles was not caused by different sensitivities to GdA in different strains, we assayed the Hsp90 activity by measuring the kinase activity of v-Src, a well-known Hsp90 client (Xu and Lindquist 1993), with or without GdA treatments (Supplemental Fig. S5A and Methods). The v-Src kinase activity was reduced to the background level in all tested strains upon GdA treatment, indicating that the sensitivity difference to GdA is unlikely the primary reason for the observed expression differences.

We also noticed that the ML strain had a longer doubling time than the other strains when treated with 50 μM GdA (Supplemental Fig. S5B) due to the combination of a long lag time and slow growth. It raised the possibility that the expression difference may result from the secondary effect of slow cell growth. We performed an additional RNA-seq using the ML cells treated with 25 μM GdA. Under this condition, the ML strain did not exhibit such slow growth, but its transcriptome profile is still more similar to the ML strain treated with 50 μM GdA than the other strains (Spearman’s correlation, ρ=0.83, 0.81, 0.74, 0.72, and 0.59 for 25 μM GdA-treated ML compared to 50 μM GdA-treated ML, NA, WA, SK, and LAB, respectively). Next, we compared the transcriptome profiles of 50 μM-GdA treatment with other stress conditions that slow down cell growth (Supplemental Table S2). The correlation between the GdA treatment and other stress conditions is much lower than the correlation between different GdA-treated strains. These data suggest that the observed regulatory effects are not the result of slow growth in GdA-treated cells.

Moreover, we examined the expression of several known stress response genes, including *HSP104, HSP90, HSP70, HSP42*, and *HSF1* (Supplemental Table S3) (Dittmar and Pratt 1997; Glover and Lindquist 1998; Kempf et al. 2017; Masser et al. 2019)}. The mRNA levels of Hsp90 (encoded by *HSC82* and *HSP82*) and its cochaperone Hsp70 (encoded by *SSA1, SSA2, SSA3*, and *SSA4*) were induced in multiple strains probably due to the compensation effect of Hsp90 inhibition (Clarke et al. 2000; Maloney et al. 2007). On the other hand, *HSP104, HSP42*, and *HSF1* were not induced, suggesting that the GdA treatment did not trigger a general stress response. Together, our data indicate that differential gene expression observed in the GdA-treated cells is specific to Hsp90 inhibition.

To investigate if these Hsp90-dependent genes have particular biological functions and if individual strains exhibit differentially-responsive gene sets, we further investigated gene ontology (GO) for biological processes among impacted genes. Among the 26 enriched GO terms that we identified, only five were shared by more than three strains (Fig. 1A), with these latter likely representing the general physiological effects of Hsp90 or ancestral cryptic genetic variation shared by multiple strains. For instance, Hsp90 and its co-chaperones have been shown to influence mitochondrial protein import, with this function being conserved in both yeast and mammalian cells (Young et al. 2003; Hoseini et al. 2016), perhaps explaining GO enrichment for cellular respiration and response to oxidative stress. Nevertheless, even for commonly shared GO terms, we observed different levels of enrichment among strains (Fig. 1). These data indicate that the general effect of Hsp90 on individual cellular processes can be further adjusted depending on genetic background.

**Figure 1.**
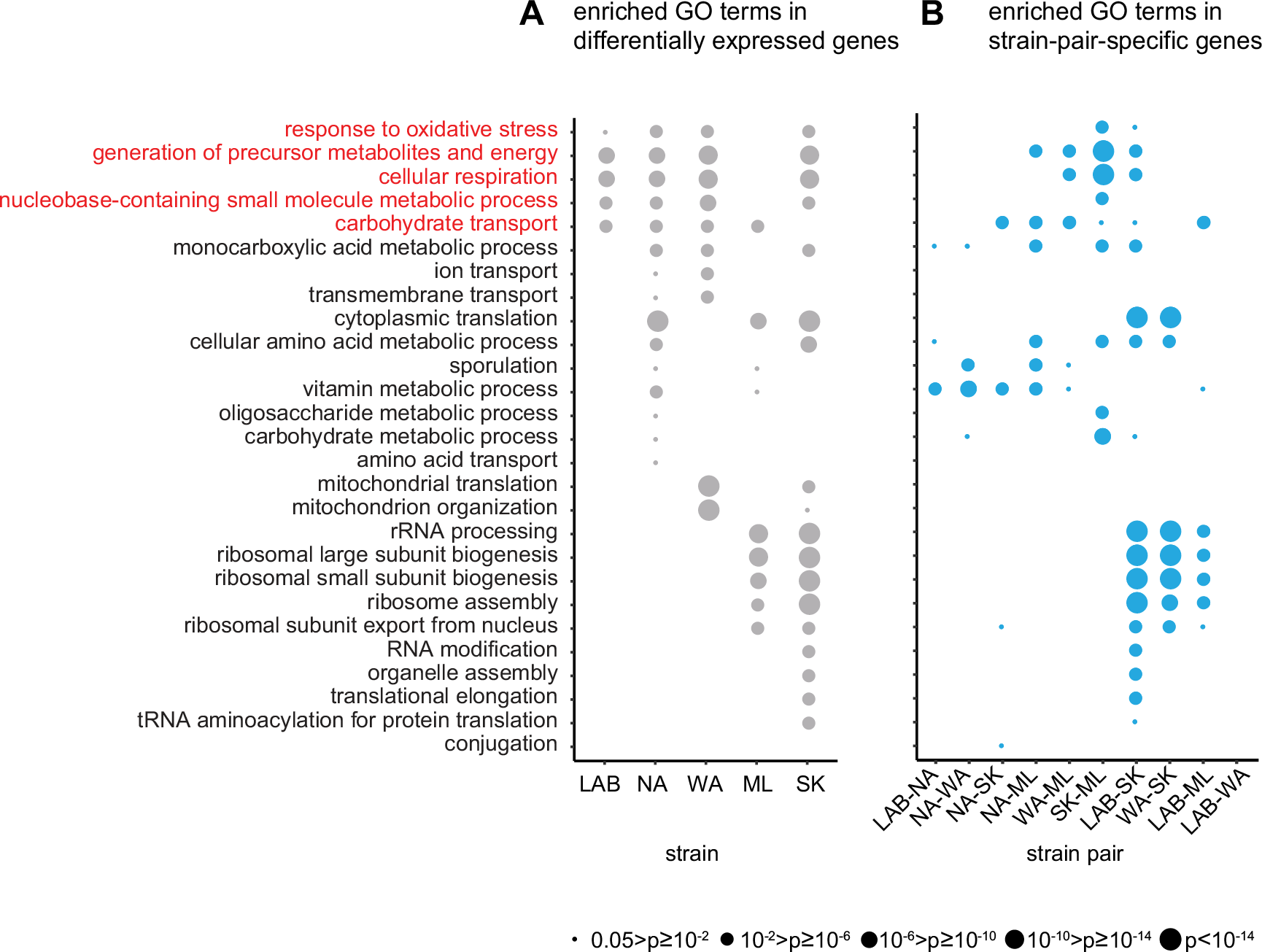
Hsp90-dependent transcriptome analysis of five diverse yeast strains. (A) GO analysis of genes differentially expressed in response to Hsp90 inhibition (+GdA/−GdA) for each strain. Only enriched GO terms have been plotted and circle size represents adjusted p-values. GO terms enriched in more than three strains were classified as a common response GO term (shown in red). (B) GO analysis of the strain-pair-specific genes for all strain pairs. Those genes exhibited more than a two-fold difference in Hsp90-dependent expression between two strains (see Supplemental Fig. S3).

In addition to common Hsp90 responses, we observed many strain-specific responses that were only enriched in one or two strains (Fig. 1A), suggesting that different strains can display unique Hsp90-dependent expression patterns. We used them to further dissect the underlying mechanisms and possible physiological effects.

### Hsp90-buffering events occur more frequently than potentiated events in yeast populations

Since we had already observed many strain-specific responses in gene expression, next we searched for differentially expressed genes that exhibited apparent differences in Hsp90 dependency between strain pairs. To do this, we compared fold-changes in mRNA abundance in cells of two given strains subjected to either of two conditions (i.e., GdA treatment or lacking GdA treatment). If a gene presented a between-strain ratio greater than 2 or less than 0.5, that gene was defined as a strain-pair-specific gene (Supplemental Fig. S3 and Table S4). In later experiments, we used this strain-pair-specific gene information to facilitate the identification of the TFs that are influenced by Hsp90 in a strain-specific manner.

When we performed GO enrichment analysis on strain-pair-specific genes, we found that the enriched GO terms strongly overlapped with those from differentially expressed genes (Fig. 1). This outcome indicates that many differentially expressed genes indeed exhibit significant differences in their Hsp90-dependencies among strains. We further divided the identified strain-pair-specific genes into buffered and potentiated groups depending on whether the differential expression between two strains under GdA-treated conditions was greater or smaller than that under untreated conditions, respectively (Supplemental Fig. S3 and Table S4). We identified more buffered genes than potentiated genes (on average 59.3% strain-pair-specific genes are buffered, one-sample t-test, p=0.02) (Supplemental Fig. S6 and Table S5), which is consistent with the hypothesis that buffered mutations are more likely to be neutral and accumulate during evolution (Geiler-Samerotte et al. 2016). In contrast, potentiated mutations are the outcome of positive selection and probably represent rare events.

Genetic buffering is known to increase the complexity of the phenotype (Sangster et al. 2004); (Jarosz et al. 2010; Lehner 2011). Using the transcriptome profiles to construct expression distance trees (see Methods), it shows that the expression tree in the normal condition (−GdA) is highly similar to the genetic distance tree (Spearman’s correlation, ρ=0.71, Supplemental Fig. S1A and S1B). Again, the LAB and ML strains show the longest distance. The NA and SK strains have the second smallest distance (that is slightly longer than the distance between NA and WA). On the other hand, the expression tree in the Hsp90-inhibition condition (+GdA) is very different from the genetic distance tree (Spearman’s correlation, ρ=−0.11, Supplemental Fig. S1C). These data suggest that the transcriptomic profiles diverge over time, similar to the genetic distance. However, the Hsp90 inhibition adds another layer of complexity into the divergence as predicted by the capacitor hypothesis.

### Identification of TFs that contribute to variable Hsp90-dependent gene expression

One of the major regulators of gene expression are TFs, which recognize specific DNA sequences (also called transcription factor binding sites or TFBSs) within promoter regions and then activate/suppress expression of the target genes (Hahn and Young 2011). Since genes involved in similar cellular processes may be regulated by the same TFs, we decided to examine if the strain-specific effects of Hsp90 on differential gene expression are mediated through specific TFs.

First, we analyzed the distribution of TFBSs among different strains. Although information on TFBS location from chromatin immunoprecipitation (ChIP)-on-chip or ChIP-seq experiments are widely used in analyses of TF-based regulation, such experiments in yeast have typically been done under normal growth conditions and in laboratory strains (Lee et al. 2002; Harbison et al. 2004; Venters et al. 2011). Inhibition of Hsp90 triggers physiological changes and leads to altered transcriptional regulation (Gopinath et al. 2014). Moreover, several studies have shown that TF-bound targets can be dramatically different among strains and that these differences in TF binding are often associated with levels of sequence variation (Zheng et al. 2010; Gallagher et al. 2014). Thus, current experimentally-validated TFBS data do not provide sufficient information to reveal the impacts of Hsp90 inhibition or how they differ among different strains. To gain a global picture of how TFBSs are distributed in our selected strains, we used a bioinformatics pipeline to predict potential TFBS via a genome-wide promoter scan based on known binding motifs (Supplemental Fig. S7, also see Methods) (Spivak and Stormo 2012). This analysis pipeline was highly sensitive and, inevitably, generated some false-positive signals. However, we found that this issue was not a serious concern since many of our predicted results could be validated by further experiments (see later sections). Among all predicted TFBSs, only the ones that were conserved across strains (96%) were used for further analyses (Supplemental Table S6).

After constructing the TFBS map, we grouped genes sharing the same TFs, and then searched for TFs whose downstream targets exhibited differential Hsp90-dependent expression among strains. We hypothesized that if a TF or its upstream signal transduction components contain strain-specific genetic variations that are buffered or potentiated by Hsp90, such events may be revealed in the expression profiles of the TF target genes when Hsp90 is compromised. Among 187 analyzed TFs, we identified 21 TFs whose targets showed significant differences in expression profile among our five tested strains (Supplemental Fig. S7 and S8, Table S7). Interestingly, Hsp90-interacting TFs are significantly enriched in our candidate list (7 out of 21 in our list versus 23 out of 187 in the whole genome, hypergeometric test, p = 0.0011). Previously, our identification of strain-pair-specific genes allowed us to detect the differentially expressed genes that exhibited strong strain-specific Hsp90-dependencies (Fig. 1, Supplemental Fig. S3, S6, Table S4, and S5). We used this information as another criterion to further refine our list of TF candidates. If strain-pair-specific genes are enriched among the targets of a specific TF, it further supports that this TF may have strong impacts on strain-specific differential gene expression. Eleven TFs were observed to exhibit such enrichment in at least one strain pair (Fig. 2, Supplemental Table S7, and S8). Moreover, some of those TFs shared similar enrichment patterns (Fig. 2, Supplemental Fig. S9). Next, we selected six of the eleven TFs (Msn2, Msn4, Com2, Rap1, Dot6, and Tod6), representing different expression patterns among strain pairs, for experimental validation.

**Figure 2.**
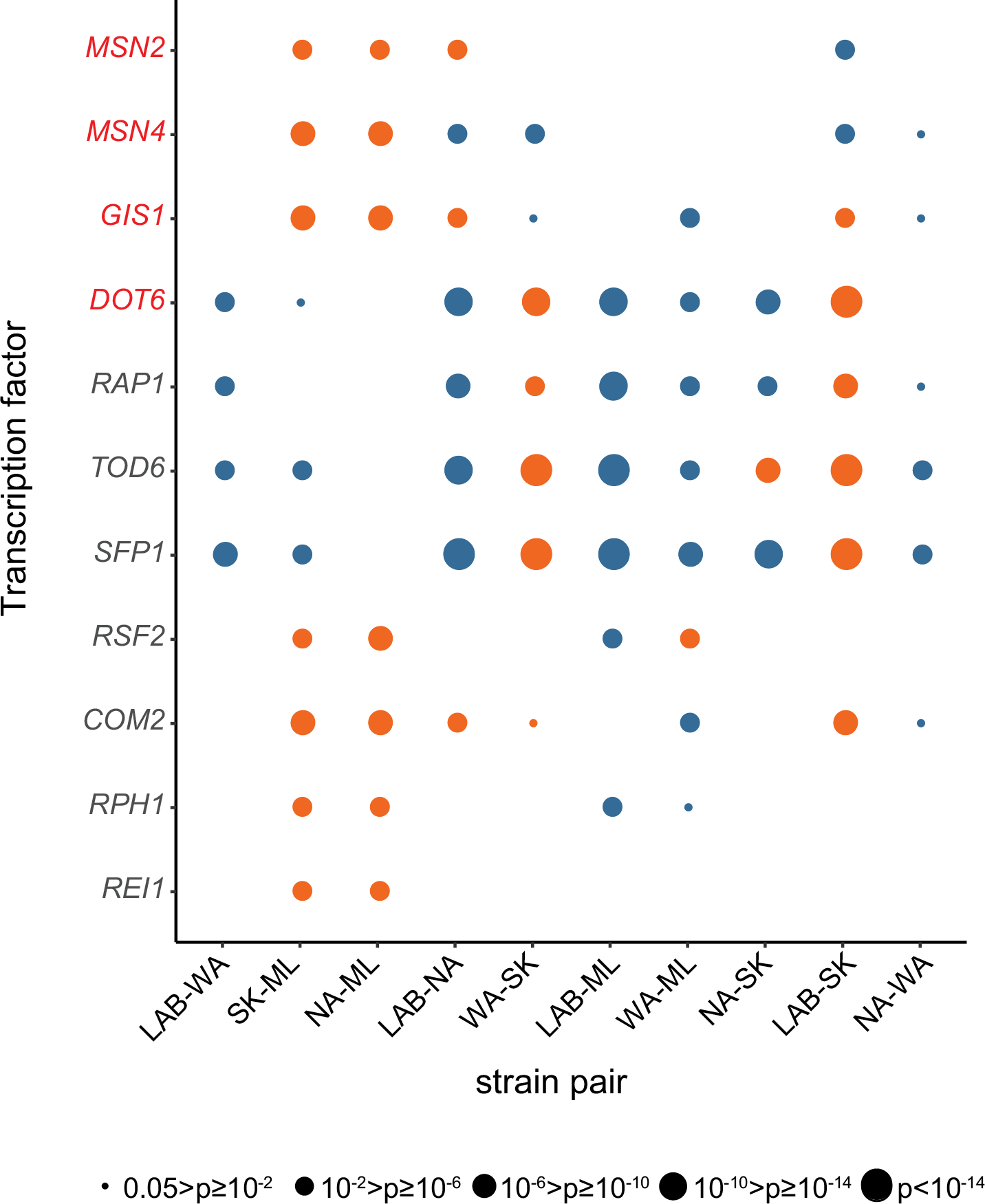
Candidate transcription factors involved in Hsp90-dependent strain-specific regulation. The targets of these candidate TFs show significant differences in between-strain Hsp90-dependent expression. Dot size represents Bonferroni-adjusted p-value from the Wilcoxon rank-sum test (Table S7). Orange dots represent those strain-pair-specific genes that are also enriched among TF targets (Table S8). TF names are labeled in red on the Y-axis if they are Hsp90 clients.

### Hsp90 inhibition leads to strain-specific upregulation of Msn4

Msn2 and Msn4 physically interact with Hsp90 (Truman et al. 2015), and they are functionally redundant paralogs that activate genes involved in stress responses (Estruch and Carlson 1993; Morano et al. 2012). We found that when Hsp90 was compromised, target genes of Msn2 and Msn4 in the SK and NA strains were significantly upregulated relative to in the ML strain (Fig. 3A). However, since both Msn2 and Msn4 can bind to the same stress-responsive element (a DNA motif containing AGGGG) and they share a large proportion of their targets (Wieser et al. 1991), it raises the possibility that only one of them is responsible for the observed changes in expression. The identification of both TFs as Hsp90-dependent TFs may be due to the shared targets and it is hard to distinguish this possibility by computational analysis.

**Figure 3.**
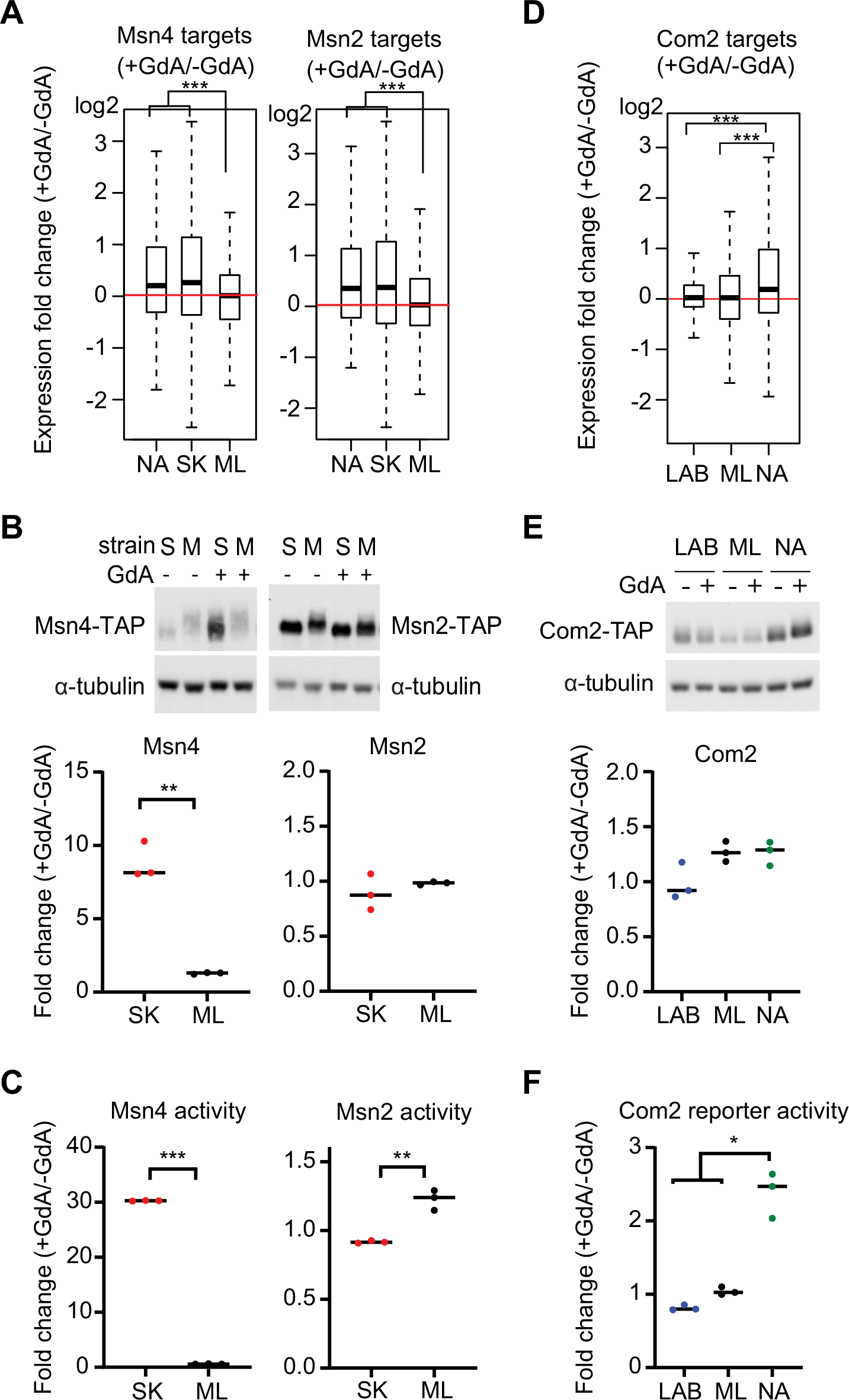
The impact of Hsp90 on Msn4 and Com2 varies among strains. (A) Expression profiles of Msn4 and Msn2 targets reveal significant differences in Hsp90 dependency between the SK and ML strains (two-sided Wilcoxon rank-sum test) (Table S8). The medium values are shown by the horizontal line in the box plots. The whiskers above and below the box indicate the 1.5x interquartile range. The outliers are labeled if the value is out of the 1.5x interquartile range. (B) Msn4 is highly induced in the SK strain (S), but not in the ML strain (M), upon Hsp90 inhibition (two-sided unpaired t-test with Welch’s correction, p = 0.0091 for Msn4, p = 0.4504 for Msn2). Msn2- and Msn4-TAP fusion protein levels from the SK and ML strains were measured by Western blot. Samples were cultured in the absence (-) or presence (+) of the Hsp90 inhibitor GdA. α-Tubulin is an internal control. Quantification of the Western blot results is shown in the bottom panel. (C) Msn4 activity is strongly enhanced in the SK strain, but not in the ML strain, upon Hsp90 inhibition (two-sided unpaired t-test with Welch’s correction, p<0.0001 for Msn4, p=0.0618 for Msn2). The transcriptional activities of Msn2/4-lexA fusion proteins were measured by one-hybrid assay in which the reporter (β-galactosidase) is transcriptionally regulated by the lexA fusion protein and β-galactosidase activity is used to represent relative TF activity. (D) Expression profiles of Com2 targets reveal significant differences in Hsp90 dependency between the LAB, ML and NA strains (two-sided Wilcoxon rank-sum test) (Table S8). The medium values are shown by the horizontal line in the box plots. The whiskers above and below the box indicate the 1.5x interquartile range. The outliers are labeled if the value is out of the 1.5x interquartile range. (E) All three strains show similar levels of altered Com2 protein abundance (two-sided unpaired student t-test with Welch’s correction, p=0.079 between LAB and ML, p=0.0845 between LAB and NA, p=0.9936 between NA and ML). Abundances of Com2-TAP protein were measured and quantified by Western blot. (F) Expression of the Com2 reporter is significantly increased in the NA strain upon Hsp90 inhibition (two-sided unpaired t-test with Welch’s correction, p=0.0121 between LAB and NA, p=0.0155 between NA and ML). Transcriptional activity of the Com2-lexA fusion protein was measured by one-hybrid assay. *, adjusted p-value < 0.05; **, adjusted p-value < 0.01; ***, adjusted p-value < 0.001.

To further establish the effect of Hsp90 on Msn2 and Msn4, we measured protein abundances of Msn2 and Msn4 in response to Hsp90 inhibition (GdA treatment). We found that, in the SK strain in which Msn2 and Msn4 targets were upregulated in Hsp90-compromised cells, Msn4 was also highly induced (Fig. 3B). In contrast, the ML strain maintained similar levels of Msn4 under GdA-treated and –untreated conditions. Moreover, Msn2 abundance was not affected by Hsp90 inhibition in either strain (Fig. 3B). Since the functions of Msn2 and Msn4 are also regulated by post-translational modifications and nuclear translocation (Gorner et al. 1998), we performed one-hybrid reporter assays to directly measure the transcriptional activities of Msn2 and Msn4 (see Methods). Consistent with our protein abundance data, Msn4 activity was increased ∼30-fold in the GdA-treated SK strain relative to that of the GdA-treated ML strain (Fig. 3C). Msn2 only showed mild Hsp90-dependent differences in activity (<1.5-fold) between strains. To ensure the observed effect is really Hsp90-dependent, we repeated the experiment using a different Hsp90 inhibitor, radicicol, and observed a similar SK-specific Msn4 induction (Supplemental Fig. S10A). Together, these data suggest that Msn4 is the dominant factor contributing to the differential Hsp90-dependent expression of Msn4 targets.

### Hsp90 affects Com2 activity without changing its protein abundance in the NA strain

Com2 shares a similar core motif (AGGGG) with Msn2 and Msn4 (Supplemental Table S7), but its function is not well characterized (Spivak and Stormo 2012; Kahana-Edwin et al. 2013). In our reporter enrichment analysis, Com2 exhibited a different pattern to that of Msn2/4 (Fig. 2), suggesting that Com2 targets were affected by Hsp90 via different regulatory mechanisms. Basal protein levels of Com2 varied among strains (Fig. 3E), but the alterations in protein abundance due to Hsp90 inhibition did not differ significantly among strains. This outcome indicates that specific upregulation of Com2 targets in the NA strain (Fig. 3D) was probably not mediated by altered Com2 abundance (Fig. 3E). However, Com2 transcriptional activity of the NA strain, but not the LAB or ML strains, increased significantly under GdA treatment (Fig. 3F). Thus, post-translational regulation is probably involved in the Hsp90-dependent effect of Com2 observed here, unlike the case of Msn4.

### Hsp90 influences the expression of ribosomal genes through multiple TFs

Since we noted that several enriched GO terms of the differentially expressed genes were related to ribosomal function (Fig. 1), we next focused on Rap1, an essential TF involved in diverse cellular processes, including chromatin silencing, telomere length maintenance and transcriptional activation of ribosomal genes (Bosio et al. 2011; Azad and Tomar 2016)}. Rap1 alters nucleosome position and recruits the cofactors Fhl1, Ifh1, Sfp1, and Hom1 to regulate ribosomal genes (Reja et al. 2015). Interestingly, Rap1 targets in the SK, WA, and LAB strains were differentially impacted upon Hsp90 inhibition (Fig. 4A).

**Figure 4.**
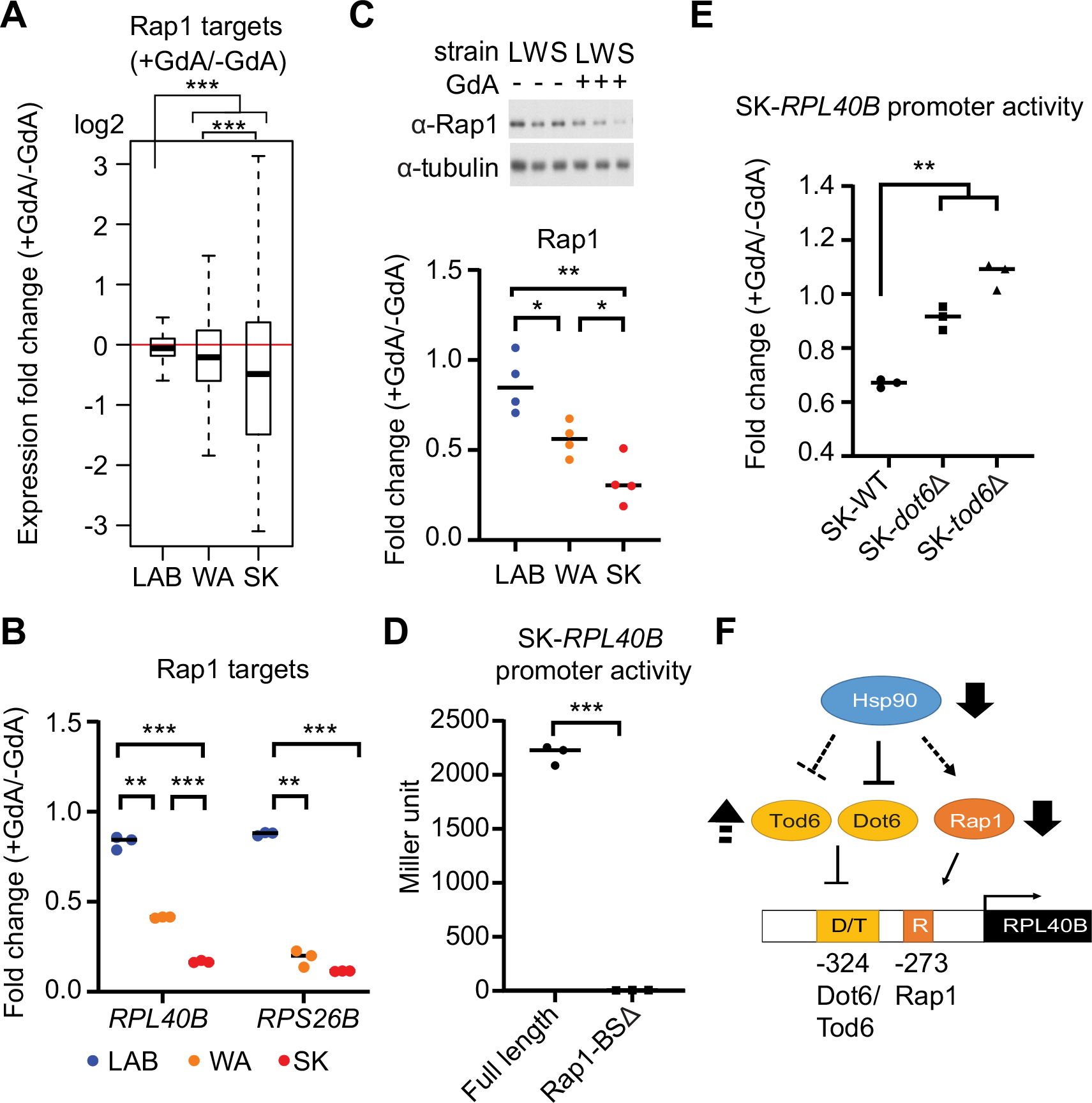
Hsp90 regulates the expression of ribosomal genes through a complex regulatory network. (A) Expression profiles of Rap1 targets are significantly different among the LAB, WA, and SK strains (Wilcoxon rank-sum test) (Table S8). The medium values are shown by the horizontal line in the box plots. The whiskers above and below the box indicate the 1.5x interquartile range. The outliers are labeled if the value is out of the 1.5x interquartile range. (B) Two Rap1-regulated ribosomal genes, *RPL40B* and *RPS26B*, exhibit strong, intermediate or weak Hsp90-dependent effects in the SK (S), WA (W), and LAB (L) strains, respectively (two-sided unpaired t-test with Welch’s correction, p=0.0024, 0.0008, and <0.0001 for *RPL40B* in pair LAB-WA, LAB-SK, and WA-SK, respectively; p=0.001, <0.0001, and 0.1114 for *RPS26B* in pair LAB-WA, LAB-SK, and WA-SK, respectively). Fold-change in expression (+GdA/−GdA) of *RPL40B* and *RPS26B* was measured by real-time PCR. (C) The protein abundance of Rap1 is reduced to different extents upon Hsp90 inhibition (two-sided unpaired t-test with Welch’s correction, p=0.0235, 0.0024, 0.0326 in pair LAB-WA, LAB-SK, and WA-SK, respectively). Rap1 protein level was measured by Western blot. (D) Rap1 is a crucial transcriptional activator for expression of *RPL40B*. The activities of full-length and Rap1 binding site-deleted (Rap1-BSΔ) SK-*RPL40B* promoters were tested in the SK strain. Removal of the Rap1 binding site led to a strong reduction in gene expression (two-sided unpaired t-test with Welch’s correction, p=0.0006). (E) Dot6 and Tod6 are involved in Hsp90-dependent regulation of expression in the SK strain. Promoter activity became almost Hsp90-independent (fold change was approached to 1) when either *DOT6* or *TOD6* was deleted from the SK strain (two-sided unpaired t-test with Welch’s correction, p=0.0056 between wild type and *dot6Δ* mutant, p=0.0026 between wild type and *tod6Δ* mutant). (F) A model describing how Hsp90 influences the expression of ribosomal genes via Rap1, Dot6, and Tod6. The dashed lines represent the regulation indirectly inferred from genetic data. *, adjusted p-value < 0.05; **, adjusted p-value < 0.01; ***, adjusted p-value < 0.001.

We examined two known Rap1-regulated ribosomal genes, *RPL40B* and *RPS26B*, using quantitative PCR (q-PCR) to determine if differential expression patterns were similar for individual genes (Supplemental Table S9). Consistent with the overall Rap1 target data (Fig. 4A), both of these genes exhibited strong, intermediate, or weak Hsp90-dependent effects in the SK, WA, and LAB strains, respectively (Fig. 4B, Supplemental Fig. S10B). A similar pattern of Hsp90 dependency was also observed for Rap1 protein abundance (Fig. 4C). Together, these results indicate that Hsp90 can influence the same TF to different extents depending on strain background. We also constructed an SK-*RPL40B* promoter-lacZ reporter gene to investigate to what extent Rap1 contributes to *RPL40B* expression. When we deleted the Rap1 binding site (CCATCCGTGCCTC; located 273 base pairs upstream of the translation start site) from the SK-*RPL40B* promoter, we found that β-galactosidase activity was drastically reduced in the SK strain (Fig. 4D), confirming that Rap1 is a crucial activator of ribosomal gene expression.

Our list of six TFs contains a pair of paralogs, Dot6 and Tod6, which inhibit ribosomal gene expression in response to environmental change (Lippman and Broach 2009). As for Msn2 and Msn4, Dot6 and Tod6 bind to similar binding motifs and share a significant proportion of their targets. However, Dot6 and Tod6 are controlled separately by the TORC1 and PKA pathways, respectively, indicating that these two TFs have diverged and are controlled by different upstream signals in response to diverse environments (Lippman and Broach 2009). Besides, Hsp90 physically interacts with Dot6 but not Tod6 (Oughtred et al. 2016). To investigate the contribution of Dot6 and Tod6 to differential Hsp90-dependent gene expression, we performed promoter activity assays on the SK-*RPL40B* promoter-lacZ reporter in the SK strain. Our TFBS analysis indicated that the *RPL40B* promoter contains both Rap1 and Dot6/Tod6 binding sites (Supplemental Table S7). We used a reporter construct lacking the Rap1 binding site to directly measure the effects of Dot6 and Tod6. When we deleted *DOT6* or *TOD6*, expression of the SK-*RPL40B* promoter-lacZ reporter became almost Hsp90-independent in the SK strain (Fig. 4E). Thus, unlike Msn2 and Msn4, both Dot6 and Tod6 play a role in strain-specific Hsp90 dependency. Together, these findings reveal that differential Hsp90-dependent expression of ribosomal genes is controlled by a complex network of TFs that includes the transcription activator Rap1, as well as the repressors Dot6 and Tod6 (Fig. 4F).

### Hsp90-dependent expression profiles are correlated with cellular responses to environmental change

Our experiments validated that Hsp90 exerts differential effects on several TFs depending on the strain background, leading to strain-specific differential expression of many target genes. However, such differences will only impact population evolution if they result in physiological changes in the cell. Msn2 and Msn4 are known to be involved in oxidative stress responses (Morano et al. 2012). Since we observed Hsp90-dependent upregulation of Msn4 in the SK strain (Fig. 3), we tested if such a change could lead to fitness differences between the SK and ML strains under oxidative stress. In the absence of H_2_O_2_, inhibition of Hsp90 had only a mild effect on growth of the ML and SK strains. However, when we challenged cells with H_2_O_2_, the ML strain displayed greater sensitivity to that oxidative stress than the SK strain (Fig. 5A).

**Figure 5.**
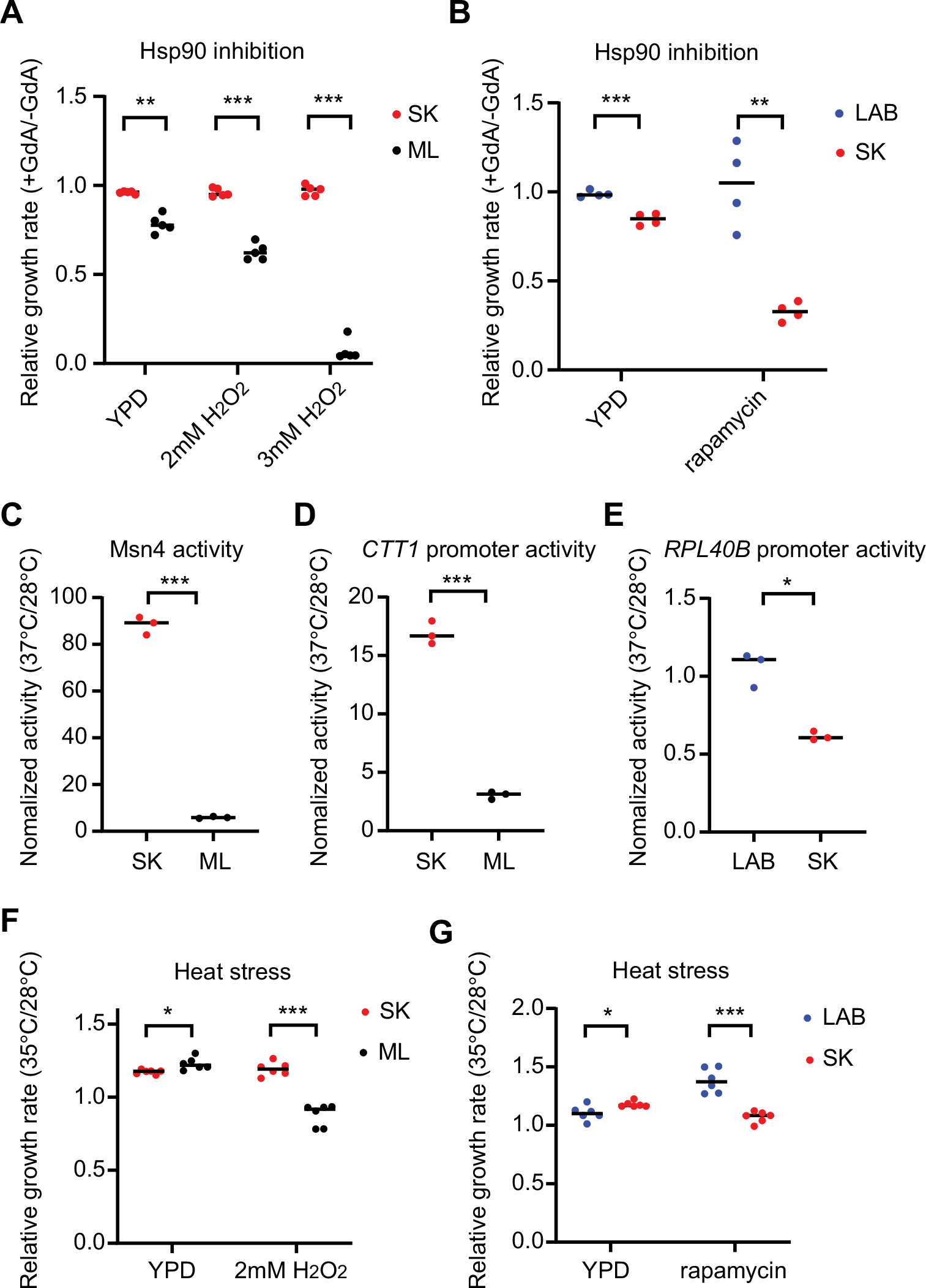
Hsp90-dependent expression profiles are correlated with cellular responses to stress and heat stress triggers yeast cells to manifest Hsp90-dependent strain-specific phenotypic variation. (A) The ML strain is more sensitive to the oxidative stress induced by H_2_O_2_ treatments than the SK strain when Hsp90 is compromised (two-sided unpaired t-test with Welch’s correction between strains, p=0.0012 for YPD condition, p<0.0001 for oxidative conditions). This result is consistent with the observation that Msn4 is highly induced by Hsp90 inhibition only in the SK strain. (B) The LAB strain is more resistant to the TORC1 inhibitor, rapamycin, than the SK strain upon Hsp90 inhibition (two-sided unpaired t-test with Welch’s correction between strains, p=0.0008 for YPD, p=0.0075 for the rapamycin condition). This result is consistent with the observation that the SK strain shows a stronger Hsp90-dependent reduction in expression of ribosomal genes than the LAB strain. (C) Msn4 activity is induced under heat stress in the SK strain, but not in the ML stain (two-sided unpaired t-test with Welch’s correction between strains, p=0.0006). Msn4 activity was measured by one-hybrid assay before and after heat shock (37 ℃) for 1 h. (D) Expression of *CTT1*, a target of Msn4, shows an induction pattern similar to that of Msn4 activity after heat shock (37 ℃) for 1 h (two-sided unpaired t-test with Welch’s correction between strains, p=0.0007). (E) Expression of *RPL40B* is reduced more in the SK strain than in the LAB strain under heat stress (two-sided unpaired t-test with Welch’s correction between strains, p=0.0162). β-Galactosidase assays were performed before and after heat shock for 1 h using the strains carrying the *RPL40B* promoter-lacZ reporter. (F) The SK strain grows better than the ML strain in the presence of oxidative stress at high temperatures (two-sided unpaired t-test with Welch’s correction between strains, p=0.0205 for YPD, p<0.0001 for the H_2_O_2_ condition). (G) The LAB strain grows better than the SK strain in the presence of rapamycin at high temperatures (two-sided unpaired t-test with Welch’s correction between strains, p=0.0319 for YPD, p=0.0003 for the rapamycin condition). In F & G, we chose 35 ℃ for high-temperature growth since some tested strains grew very slowly at 37 ℃. *, adjusted p-value < 0.05; **, adjusted p-value < 0.01; adjusted p-value < 0.001.

Given that expression of ribosomal genes controlled by Rap1, Dot6, and Tod6 also exhibited different levels of Hsp90 dependency between the LAB and SK strains (Fig. 4), we directly tested if these two strains had different resistances to rapamycin, a drug that represses the expression of ribosomal genes by inhibiting the TOR pathway (Pestov and Shcherbik 2012). Consistent with levels of Hsp90 dependency among its impacted genes, the SK strain indeed showed lower resistance to rapamycin when Hsp90 was compromised (Fig. 5B). Our results demonstrate a direct link between strain-specific differential expression and altered cell physiology, suggesting that the observed buffering or potentiating effects of Hsp90 have the potential to influence population adaptability.

### Heat stress triggers yeast cells to manifest strain-specific phenotypic variation

The capacitor hypothesis predicts that cryptic genetic variation buffered by Hsp90 allows cells to manifest phenotypic variation when faced with environmental challenges (Sangster et al. 2004). We already revealed several strain-specific phenotypic variations upon inhibiting Hsp90 by GdA treatment. Could such variations also be manifested when cells encounter environmental stress, as predicted by the capacitor hypothesis? Previous studies have suggested that the massive amounts of misfolded proteins induced by acute environmental stress could titrate out the cytosolic activity of Hsp90 and cause Hsp90-compromised phenotypes (Rutherford and Lindquist 1998; Hsieh et al. 2013; Karras et al. 2017). To test this possibility, we challenged different yeast strains with heat stress to determine if we could detect strain-specific phenotypic variation as observed under GdA-treated conditions.

First, we measured Msn4 transcriptional activity via one-hybrid reporter assays with or without heat stress. Although heat stress induces Msn4 activation (Martinez-Pastor et al. 1996), we observed much higher induction in the SK strain compared to in the ML strain (Fig. 5C). We also measured the expression of an Msn4 target, *CTT1* (Supplemental Table S9), using the *CTT1* promoter-lacZ reporter. Consistent with the Msn4 activity data, *CTT1* expression was much higher in the SK strain upon heat shock (Fig. 5D). Moreover, when we deleted the Msn4 binding site in the *CTT1* promoter (AAGGGGC; located 364 base pairs upstream of the translation start site), the induction of *CTT1* was significantly reduced in the SK strain but not in the ML strain (Supplemental Fig. S10C). These data provided evidence that Msn4 was the major contributor to the heat-induced strain-specific induction of *CTT1*. We also used the *RPL40B* promoter-lacZ reporter to investigate how heat stress affected the expression of ribosomal genes. Similar to GdA treatment (Fig. 4E), expression of the *RPL40B* promoter was significantly reduced only in the SK strain after heat stress (Fig. 5E).

Through our GdA-treatment experiments, we showed that the observed strain-specific patterns of differential expression are correlated with fitness differences under changing environments (Fig. 5A and 5B). Extending that analysis, we challenged cells under a heat stress environment with the additional challenge of either oxidative stress or rapamycin treatment. Similar to our findings from GdA treatment, under heat stress the ML strain exhibited greater sensitivity to H_2_O_2_, whereas the LAB strain grew better than the SK strain in the presence of rapamycin (Fig. 5F and 5G). Together with our gene expression data, our results show that heat stress can trigger cells to manifest strain-specific phenotypic variation, just as treatment with a specific Hsp90 inhibitor can. Such buffered variation allows cell populations to maintain a stable phenotype under normal conditions but to manifest diverse phenotypes when they encounter stressful scenarios.

## Discussion

Buffering or potentiation of genetic variation by Hsp90 has been suggested to influence organism evolution (Sangster et al. 2004; Jarosz et al. 2010). In recent years, Hsp90-buffered or -potentiated events have been further implicated in human diseases and cancer development (Jarosz 2016; Karras et al. 2017). However, detecting such genetic variation is challenging, despite the molecular and genetic properties of Hsp90 having been investigated thoroughly (Taipale et al. 2010; Schopf et al. 2017). By analyzing differential gene expression patterns among five diverse yeast strains, we identified 11 TFs that are subject to Hsp90-dependent and strain-specific regulation (Fig. 2). We performed experiments to further confirm the involvement of five TFs in the observed patterns of gene expression and predicted impacts on cellular physiology. Together with various enriched GO terms observed in differentially expressed genes, our data suggest that strain-specific Hsp90-dependent regulation is prevalent in natural yeast populations.

Hsp90 may regulate cryptic genetic variation through two different mechanisms (Rutherford et al. 2007). In the direct buffering (or potentiating) model, the cryptic variation resides in the Hsp90 client. Hsp90 directly interacts with the variant-carrying protein to maintain its function or protein stability (Fig. 6). Some disease-related mutations and oncogene mutants likely belong to this category (Blagosklonny et al. 1995; An et al. 2000; Karras et al. 2017). The indirect buffering (or potentiating) model suggests that the variant-carrying protein is not itself an Hsp90 client, but interacts with the client protein or pathway. When Hsp90 is inhibited, the compromised client synergistically enhances the mutant effect, leading to a detectable phenotype (Fig. 6). Hsp90-buffered gene expression arising from repression of endogenous retroviruses in mice represents a well-documented case of indirect buffering (Hummel et al. 2017). Although Hsp90 clients only represent 8.8% of the yeast proteome (Oughtred et al. 2016), Hsp90 can impact a much broader range of proteins through indirect buffering (or potentiation).

**Figure 6.**
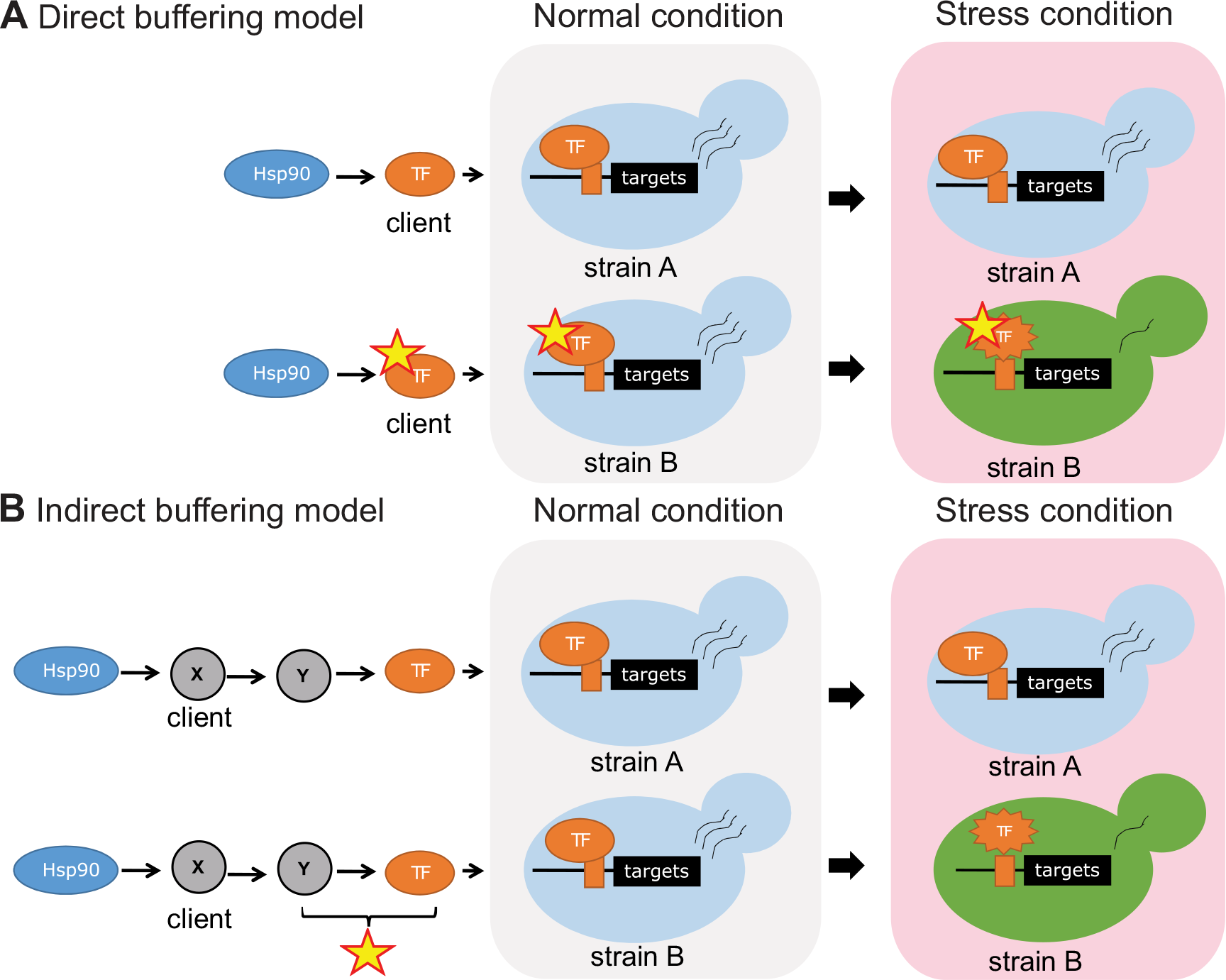
Two models showing how Hsp90 can buffer a variant-carrying protein in a transcriptional regulatory pathway. (A) In the direct buffering model, the variant-carrying TF is an Hsp90 client and its function is stabilized by Hsp90. When the cell encounters stress, Hsp90 is recruited to clear misfolded proteins caused by stress, so the TF function is no longer buffered, allowing differential gene expression. (B) In the indirect buffering model, the TF function is regulated by protein X, which is an Hsp90 client. When X is fully functional, the variant-carrying TF or protein Y does not show obvious defects. However, when X is compromised after titration of Hsp90 (by stress-induced misfolded proteins), the combined effects of compromised X and variant-carrying TF (or Y) result in differential gene expression.

Seven out of the 11 high-confident Hsp90-dependent TFs do not directly interact with Hsp90. These non-client TFs can be regulated by several possible mechanisms. First, the non-client TFs may be the direct targets of the Hsp90-dependent client TFs. By examining the promoter regions of the seven non-client TFs, we identified the binding motifs of the Hsp90 client TFs (*DOT6, MSN4*, and *GIS1*) in five of the non-client TFs (Supplemental Table S9). However, the general expression patterns of the client TF targets are not consistent with that of the motif-containing non-client TFs and their targets except for the *DOT6*-*TOD6* pair (Fig. 2, Supplemental Fig. S9 and Table S10). Since Dot6 and Tod6 share many common targets, it remains unclear how significant such regulation is in this case. Second, the non-client TFs may be regulated by Hsp90 clients post-transcriptionally. As revealed by the phosphoGRID database (Sadowski et al. 2013), all of our non-client TFs contain multiple post-translational modification sites although the modification enzymes are not identified in most cases (Supplemental Table S10). Many protein modifiers are known to be Hsp90 clients and their activities are controlled by Hsp90 (Taipale et al. 2010; Schopf et al. 2017). In addition, our results have shown that one of our Hsp90-dependent TFs, Com2, is probably regulated by post-translational modifications (Fig. 3). We examined the one-degree interactors of the non-client TFs and identified 40 Hsp90 client proteins, including TF cofactors, Hsp90 cochaperones, protein modifiers, and ribosomal subunits (Supplemental Table S10). Hsp90 may regulate the non-client TFs through these interacting Hsp90 client proteins.

Recent studies evidence that transcriptional rewiring is quite common during yeast evolution (Dalal and Johnson 2017; Johnson 2017). Interestingly, several of our identified TFs were involved in these rewiring events. For instance, regulation of ribosomal genes was dramatically rewired from Rap1 in *S. cerevisiae* to Tbf1 in *C. albicans* (Lavoie et al. 2010; Sarda and Hannenhalli 2015). Moreover, it has been suggested that Dot6 and Tod6 targeting was rewired among yeast species based on motif changes (Sorrells et al. 2018). With regard to the regulatory pathway for responses to oxidative stress, the requisite TFs were rewired from Msn2/Msn4 in *S. cerevisiae* to Fkh2 in *C. albicans* (Sarda and Hannenhalli 2015). It has been suggested that transcriptional rewiring often occurs through an intermediate state in which regulatory changes are selectively neutral to the cells (Dalal and Johnson 2017). Since Hsp90-buffered genetic variation allows cells to accumulate neutral regulatory changes, it raises the interesting possibility that Hsp90 may play a role in facilitating transcriptional rewiring.

Altered transcriptional regulation is thought to play a crucial role in the adaptive evolution of various organisms (Carroll 2000; Wagner and Lynch 2008; Romero et al. 2012). For example, a comparison of humans with other primates has revealed that TFs are enriched among genes exhibiting a human-specific increase in expression (Gilad et al. 2006). In threespine stickleback fish, the adaptive pelvic reduction observed in different natural populations is caused by recurrent mutations in *Pitx1*, a TF required for normal hindlimb development in vertebrates (Chan et al. 2010). Repeated mutations in another TF, *optix*, were also found to be responsible for convergent evolution of wing pattern mimicry among different butterfly species (Reed et al. 2011). In plants, genetic variations in two MADS-box TFs, *SOC1* and *FUL*, alter plant growth form (annual or perennial) and longevity (Melzer et al. 2008). Adaptive evolution mediated by TF variation is equally common in microorganisms. In yeast, altered expression or activity of TFs has been shown to contribute to population- or species-level differences in morphology, environmental responses, or lifecycle (Zordan et al. 2006; Chang et al. 2008; Gerke et al. 2009; Fraser et al. 2010; Lewis et al. 2010; Engle and Fay 2012; Nagy et al. 2014; Sherwood et al. 2014). Most of the observed changes in TFs and gene expression we report in our study are probably not the outcome of adaptive evolution. However, we have shown that many Hsp90-dependent strain variations can lead to detectable phenotypic differences when the function of Hsp90 is perturbed by stress. These Hsp90-linked variations provide a large repertoire of material on which natural selection can act when cells are challenged by changing environments.

## Methods

### Yeast culture conditions

To extract total RNA or perform the TF-activity assay, yeast cells were cultured for 8 h in 2x complete supplement medium (2x CSM: 0.7% yeast nitrogen base without amino acids, 2% dextrose, 0.2% CSM powder) adjusted to pH 6 with DMSO as control or with 50 μM or 25 μM Geldanamycin (GdA) (InvivoGen, San Diego, CA, USA). Final cell density was controlled to OD_600_ = 0.2 to 0.8. The cells for promoter activity assays were cultured in cloNAT-containing medium [2x CSM supplied with 9.375 μg/ml cloNAT (Werner BioAgents, Jena, Germany)] and DMSO (D8418, Sigma-Aldrich, St. Louis, MO, USA) or 50 μM GdA. To ensure that the observed phenotypes were not GdA-specific, we also tested a different Hsp90 inhibiter radicicol (#R-1130, AG Scientific, San Diego, CA) and showed that 25 μM radicicol-treated cells exhibited phenotypes similar to GdA-treated cells (Supplemental Fig. S10A and S10B). The cells for growth curve measurement were cultured in rich medium (YPD: 1% yeast extract, 2% bactopeptone, and 2% glucose), or in YPD containing 2 or 3 mM H_2_O_2_ (#18312, Riedel-de Haen, Seelze, Germany), or in YPD containing 25 nM rapamycin (#1568, BioVision, Milpitas, CA, USA). Hsp90 inhibition was induced by treatment with 50 μM GdA.

### Yeast strain construction

All *S. cerevisiae* strains used in this study are shown in Supplemental Table S11. Primer pairs and the template DNA used for construction are listed in Supplemental Table S12. We used polymerase chain reaction (PCR) to amplify DNA fragments where necessary, and PCR products were purified using a PCR cleanup kit or gel extraction kit (K-3037, Bioneer, Oakland, CA, USA). For genome editing, DNA fragments containing the selection marker flanked by 5’ and 3’ gene-specific homologous regions were transformed into cells to replace the endogenous region through homologous recombination. For TAP tagging, the 5’ homologous region was amplified by the primers gene-F1 and gene-F1-TAP-R1, and the 3’ homologous region was amplified by the primers gene-TAP-F3 and gene-R3. The TAP and *HIS* selection markers were amplified from the TAP collection (Ghaemmaghami et al. 2003). Then, a second round of PCR was used to fuse these fragments. The final products were purified and transformed into yeasts using the lithium chloride method (Gietz and Woods 2002). Strains in which final products had been correctly integrated were selected under a selection plate and checked by PCR and sequencing.

For the one-hybrid assay, the reporter system (including lacZ driven by the promoter with eight operator sites of lexA) was derived from the pSH18-34 plasmid (Estojak et al. 1995). To increase basal promoter activity for testing repressor activity, five operator sites for Gcn4-binding were inserted downstream of the lexA operators. The entire cassette was inserted into the *HIS3* locus using the *URA*3 marker. The lexA-tagging for TFs was achieved using pSFS4A-lexA plasmid; a marker-free system derived from the pSFS2A plasmid whereby the drug-resistant gene *SAT1* can be “flipped out” by the *FLP* gene (Reuss et al. 2004). Since the *Candida albicans MAL2* promoter controlling the *FLP* gene is not well regulated in *S. cerevisiae*, we changed the *CaMAL2* promoter to the *ScGAL1* promoter using an In-Fusion kit (Takara Bio USA Inc., Mountain View, CA, USA) to generate pSFS4A plasmid. Next, the lexA-binding domain was inserted into the pSFS4A plasmid via the ApaI and XhoI sites to construct pSFS4A-lexA.

To insert lexA into the C-terminal domain of TFs, we inserted the 5’ and 3’ gene-specific homologous regions via the KpnI-ApaI and NotI-SacI sites on the pSFS4A-lexA plasmid, respectively. For the 5’ region, the homologous 300-500 bp region before the translation stop site was amplified using the primer pair TF-kpnI-F1 and TF-ApaI-R1. For the 3’ region, the homologous 300-500 bp region after the translation stop site was amplified using the primer pair TF-NotI-F3 and TF-SacI-R3. The resulting fragment was transformed into yeasts by a modified electroporation method (Hsu et al. 2011). Log-phase cells (10 ml) were harvested by centrifugation and resuspended in 10 ml lithium buffer (Tris-EDTA pH 8, 0.1 M lithium acetate pH 7.5, 25 mM dithiothreitol). The cells were cultured at 30 ℃ for 1-2 h, and then washed sequentially with 30 ml sterilized ice-cold ddH_2_O and 5 ml ice-cold 1 M sorbitol. The cells were resuspended in 500 μl ice-cold 1 M sorbitol. We then mixed 100 μl cell suspension with 1 μg DNA fragments and placed the mixture in 2 mm cuvettes. Cells were transformed using an electroporator (Gemini SC2, BTX, Holliston, MA, USA) with settings 1800 V, 200 Ω, and 25 μF. After electroporation, the cells were washed with ice-cold 1 M sorbitol and allowed to recover in YPD medium for 4-6 h at 30 ℃. Correct transformants were checked by PCR. To flip out the drug-resistant marker *SAT1*, cells were cultured in 2% galactose medium overnight to express the *FLP* gene and remove *SAT1* through recombination at the *FLP* recombination target sequence.

For the promoter activity assay, the promoter was inserted into pRS41H-lacZ plasmid using an In-Fusion kit (Takara Bio USA Inc.). The pRS41H-lacZ plasmid was constructed by inserting the lacZ fragment amplified from the pSH18-34 plasmid into the pRS41H plasmid (Estojak et al. 1995; Taxis and Knop 2006). The full-length promoters used for the activity assay were amplified using the primer pair promoter-infu-F and promoter-infu-R, and the promoter lacking TF binding sites was constructed by two-fragment PCR. Fragment I was amplified using primers promoter-infu-F and promoter-delTF-R, and fragment II was amplified using primers promoter-delTF-F and promoter-infu-R. The promoter fragment was ligated into the ApaI-linearized pRS41H-lacZ vector using an In-Fusion kit (Takara Bio USA Inc.). The sequence-checked plasmid was transformed into yeasts using the lithium chloride method (Gietz and Woods 2002).

For gene deletion, the 5’ homologous region was amplified using primers gene-pF1 and gene-del-R1, and the 3’ homologous region was amplified using primers gene-del-F2 and gene-dR2. The *URA* selection marker was amplified from the plasmid pUG72 (Gueldener et al. 2002).

### Construction of the genetic distance and expression distance trees

For calculating the genetic distance, we first downloaded the raw reads of each strain (LAB and ML strains from (Song et al. 2015), NA and SK strains from (Strope et al. 2015), and the WA strain was resequenced by this study) and mapped them to the reference genome (saccharomyces_cerevisiae_R64-2-1_20150113) by the BWA-MEM (Li and Durbin 2009). Then, we called the variants by GATK (version 3.8) and performed the hard filtering for SNPs and Indels (McKenna et al. 2010). The filtered variants were merged into the g.vcf format and then transformed into a fasta file by VCF-Kit (Cook and Andersen 2017). The genetics distance was calculated by the R package, ape (function dist.dna in the “TN93” model) (Paradis and Schliep 2019), and the neighbor-joining tree was constructed by BIONJ (Gascuel 1997). We took the average TPM value from three biological repeats divided by 1000 to calculate the euclidean distance among strains by dist function in R, and then constructed the neighbor-joining tree by BIONJ (Gascuel 1997).

### v-Src activity assay

Yeast cells were transformed with the Y316V-srcv5 plasmid so the v-Src protein abundance and activity could be measured directly (Wayne and Bolon 2007). The v-Src gene was driven by the GAL promoter. Since the GAL promoter was not induced in the WA strain in galactose-containing medium, we removed the WA strain from the assay. The cells were cultured in the CSM-URA medium with 2% glucose overnight and were refreshed in the CSM-URA medium with 2% raffinose for around 14 hours. Next, to induce the v-Src expression, log-phase cells were cultured in the CSM-URA with 2% galactose in the absence or presence of 50 μM GdA for 6 hours. The cell pellets were harvested and then total proteins were extracted for quantification by Western blotting.

### Yeast protein extraction and Western blotting

Cells were collected and resuspended in 1 ml lysis buffer (0.185 M NaOH, 0.75% β-mercaptoethanol) on ice for 15 min. To precipitate the proteins, 150 μl of 55% trichloroacetic acid (TCA) solution (T9159, Sigma-Aldrich) was added into the suspension on ice for 10 min. Then we collected the precipitated proteins by centrifugation at full speed for 10 min. The protein pellet was suspended in HU sample buffer (8 M urea, 5% SDS, 0.2 M Tris-HCl pH 6.5, 1 mM EDTA pH 8, 0.01% bromophenol blue, 120 mM Tris, 5% β-mercaptoethanol) at 65 ℃ for at least 30 min. The total protein abundance for each sample was quantified based on the total cell number measured by OD_600_. The protein was loaded into SDS-PAGE depending on the protein abundance of the targets. We loaded 0.02 OD_600_ cells for Hsp90 samples, 0.25 OD_600_ for v-Src samples, 0.5 OD_600_ for the phosphotyrosine detection, 0.2 OD_600_ for Msn2-TAP samples, 0.3 OD_600_ for Rap1 and Com2-TAP samples, and 0.65 OD_600_ for Msn4-TAP samples. Samples were separated by 7% SDS-PAGE at constant amplitude (25 mA), and transferred to Immobilon-PSQ PVDF membrane (Merck Millipore, Burlington, Massachusetts, USA) at 25V for 16 h.

The blot was blocked with 1% casein (C5890, Sigma-Aldrich) in PBST (0.1% tween20 in 1X PBS) for 1 h. Rabbit polyclonal anti-Hsp90 antibody (from Dr. Chung Wang, 1:200000 dilution) was used for Hsp90 detection. Mouse anti-V5-Tag antibody (MCA1360GA, Bio-Rad, Hercules, CA, USA) at 1:2000 dilution was used to detect V5-tagged v-Src. Mouse anti-phosphotyrosine antibody (05-321, Sigma-Aldrich) at 1:1000 was used to detect the phosphotyrosine signal. Rabbit anti-TAP antibody (CAB1001, Thermo Fisher Scientific, Waltham, MA, USA) at 1:2000 dilution was used to detect TAP-tagged Msn2, Msn4, and Com2. Mouse monoclonal anti-Rap1 antibody (sc-374297, Santa Cruz Biotechnology, Santa Cruz, CA, USA) at 1:2000 dilution was used to detect Rap1. Rabbit polyclonal anti-alpha-tubulin antibody (ab184970, Abcam, Cambridge, UK) at 1:20000 dilution was used to detect the internal control, tubulin. The blot was hybridized with antibodies for 1 h in 0.1% casein at room temperature and washed five times with PBST. Then, the blot was hybridized with ECL-anti-rabbit IgG (Jackson ImmunoResearch, West Grove, PA, USA, 1:20000 dilution) or ECL-anti-mouse IgG (Jackson ImmunoResearch, 1:20000 dilution) for 45 min and washed five times with PBST. Immuno-positive bands were visualized utilizing Western Lightning chemiluminescence reagent (PerkinElmer, Waltham, MA, USA). All the samples were measured in three or four technical repeats and quantified by the software, Image J.

### Growth rate measurement

The cells were cultured in the medium overnight. Next, we diluted the cell concentration to 2×10^5^ cells/ml. We added 120 μl for each well. OD_600_ was automatically measured by the Multiskan GO (Thermo Fisher Scientific). Due to the cell aggregation issue in some strains, we could only measure the growth rate using the non-shaking condition. However, we noticed that the ML strain had a long lag time when the concentration of GdA is 50 μM or higher. The underlying reason of the long lag remained unknown but it was not obvious when the ML cells were grown in shaking cultures.

### RNA preparation for next-generation sequencing

Overnight cell cultures were refreshed to log phase in 2x CSM medium. GdA (50 μM) or the same volume of DMSO was added and cells were grown for 8 h to reach OD_600_ = 0.3-0.7. For each strain and condition, we collected samples for three biological repeats. Total RNA was extracted using the phenol-chloroform method (Hsu et al. 2011), with some modifications. In brief, the cell pellet was suspended in 500 μl ice-cold lysis buffer (0.1 M Tris-HCl pH 7.5, 0.1 M LiCl, 2% β-mercaptoethanol, 0.01 M EDTA pH 8) and transferred into a tube containing 500 μl ice-cold PCIA (phenol:chloroform:isoamyl alcohol=25:24:1, pH 4.5) and 0.3 g of acid-washed glass beads. The cells were vortexed at maximum speed for 5 min to break cells. Then the lysates were centrifuged at full speed for 5 min at 4 ℃. We transferred the upper aqueous phase into 200 μl ice-cold PCIA, mixed it using a laboratory mixer, and then separated the aqueous and organic phases by centrifugation at full speed for 5 min at 4 ℃. We repeated this process until the interphase between the aqueous and organic phases was clean. RNA from the aqueous phase was precipitated in 70% ethanol for 2 h. The RNA precipitate was collected by full-speed centrifugation for 5 min and resuspended in nuclease-free water. Genomic DNA was removed using DNase A (TURBO DNA-free Kit, Invitrogen, Waltham, Massachusetts, USA) and RNA concentration was determined using a NanoDrop (Thermo Scientific).

Library preparation and next-generation sequencing were conducted by the Genomics Core, Institute of Molecular Biology, Academia Sinica (http://www.imb.sinica.edu.tw/mdarray/index.html). In brief, we used 2 μg total RNA for library preparation following the protocol in the TruSeq Stranded mRNA Library Prep Kit (Illumina, San Diego, CA, USA). The samples were sequenced by single-end reads (75 bp) following the protocol of the NextSeq 500 High Output kit V2 (75 cycles) (Illumina) on an Illumina NextSeq 500 system.

### Data processing for RNA sequencing

The single-end raw reads were trimmed in trimommatic v.0.36 with the parameters “ILLUMINACLIP:TruSeq3-SE.fa:2:30:10 LEADING:3 TRAILING:3 SLIDINGWINDOW:4:15 MINLEN:36” (Bolger et al. 2014). The trimmed reads were mapped to the reference transcripts (orf_coding_all.fasta, version R64-2-1) downloaded from the Saccharomyces Genome Database (SGD) using the “salmon” software in gcBias mode (Chan and Cherry 2012; Patro et al. 2017). Genes were removed from our analysis if the smallest raw read counts were fewer than 5 in all samples. Differentially expressed genes were analyzed using the R package, DESeq2 (Love et al. 2014). Genes with missing values were also removed due to the lack of p-values in differentially expressed gene analysis.

### WA strain genome sequence analysis

A high-quality genome assembly was not available for the strain we investigated from West Africa (DBVPG6044, denoted WA in this study), and the current version contains only 5452 annotated genes (Bergstrom et al. 2014). Therefore, we resequenced this strain. Genomic DNA was extracted using the phenol-chloroform method and RNA was removed by means of RNase treatment (Hsu et al. 2011). We used a Truseq DNA PCR-free LT kit (Illumina) for library preparation. The library was sequenced by Illumina NextSeq 500 with a NextSeq 500 Mid output v2 (300 cycle) sequencing kit for paired-end 150 bp read length. All sequencing protocols were carried out according to the manufacter’s instructions and using the Illumina NextSeq500 and NextSeq software (Nextseq control software NCS and Real-time Analysis software RTA). Illumina sequencing instruments generate BCL basecall files, and then we used Illumina bcl2fastq2.17 to generate fastq files and for demultiplexing. The raw reads were trimmed in trimommatic v.0.36 (Bolger et al. 2014). BWA-MEM software with default settings was used to map reads to the reference genome (saccharomyces_cerevisiae_R64-2-1_20150113) downloaded from SGD (Li and Durbin 2010; Chan and Cherry 2012). Average coverage of the WA genome is 219-fold. We called variants using GATK (version 3.8) and samtools (Li et al. 2009; McKenna et al. 2010). The variants were further filtered by coverage > 30 and allele frequency > 30%. The final variants we considered were those called by both GATK (version 3.8) and samtools. Based on these criteria, we identified 59783 variants, comprising 3448 indels and 56335 single-nucleotide mutations, compared to the reference genome (Supplemental Table S13). This result is consistent with a previous report in which the variant number ranged from 20,000-80,000 single nucleotide polymorphisms per strain (Liti et al. 2009).

### Prediction of transcription factor binding sites

The yeast genome sequences of the LAB, NA, ML, and SK strains were downloaded from different data sources (Chan and Cherry 2012; Skelly et al. 2013; Strope et al. 2015). The WA genome was resequenced in this study. The 500 bp upstream from the start codon of each coding gene was processed as a promoter sequence. We considered 5748 genes for which we had both promoter sequences and RNA expression values.

To identify the potential TFBS on the promoters of these 5748 genes, we downloaded the 191 TF binding motifs (position weight matrices (PWM)) and the recommended cutoff for defining functional TFBS from the ScerTF database (Spivak and Stormo 2012). The PWM provided the log-likelihood values of each base for each position in the motif. A motif scan was conducted repeatedly to calculate motif scores, represented by the sum of log-likelihood values of the inputted sequence upon moving one base each time along the entire promoter region. The motif scan was performed on both the forward and reverse DNA strands. Where the motif score was higher than the recommended cutoff provided by the ScerTF database, it was predicted to be a putative TFBS. This process was done using a customized python script (https://github.com/phhung1989/motif-scan). TFBS were recorded and classified (Supplemental Table S6). To classify the motifs, we first did a multiple sequence alignment for each gene. If a motif occurred within 20 bp and was the same in all strains it was classified as conserved. Motifs showing sequence variation among strains were termed conserved motifs with variations. If a motif had been gained or lost among strains, we classified it as non-conserved.

### Identification of candidate TFs

To examine if TF targets exhibited differential Hsp90-dependent expression among strains, we performed nonparametric Kruskal–Wallis tests (KW test) and then adjusted the multiple tested p-values using the Bonferroni method. Four TFs—Aca1, Cep3, Cup9, and Yrm1—were excluded from this analysis because they exhibited fewer than three targets in at least one strain. The remaining 187 TFs were tested. Twenty-one of those 187 TFs passed the KW test (adjusted p-value < 0.05). Next, we examined if expression of TF targets differed between any pair of strains by performing two-sided nonparametric Wilcoxon rank-sum tests followed by p-value adjustment using the Bonferroni method. The adjusted p-value < 0.05 was used to define significance. To further identify high-confidence candidate TFs, we tested if the strain-pair-specific genes, i.e. genes exhibiting significant differences between a given pair of strains, are enriched among TF targets using one-sided two-proportional tests (proportion greater than the background) followed by adjustment of multiple tested p-values using the Bonferroni method. Eleven high-confidence Hsp90-dependent TFs passed the test (adjusted p-value < 0.05). To cluster the TFs based on the Wilcoxon rank-sum test and enrichment test results, we labeled the TF by 0 if the adjust p-value ≥ 0.05 in the Kruskal–Wallis test for a strain pair, by 1 if the adjust p-value < 0.05 in the Kruskal–Wallis test and the adjust p-value ≥ 0.05 in the Wilcoxon rank-sum test, and by 2 if the adjust p-value < 0.05 in the Kruskal–Wallis and Wilcoxon rank-sum tests. These three groups represent non-Hsp90-dependent TFs, Hsp90-dependent TFs, and high-confident Hsp90-dependent TFs, respectively. The data were then analyzed and plotted by R. The distance matrix is calculated by the Euclidean method and the cluster method is the complete method.

### Gene ontology analysis

Biological processes as GO terms were assigned to each gene based on the go_slim_mapping.tab file downloaded from the SGD website (https://downloads.yeastgenome.org/curation/literature/). In total, 5748 genes have GO annotation, gene expression, and genomic sequence data for all analyzed strains. For each GO term, we conducted a hypergeometric test, followed by a multiple test correction using the Bonferroni method, to establish if genes are enriched for specific biological processes (adjusted p-value < 0.05).

### Quantitative real-time PCR

We subjected 2 μg total RNA to cDNA synthesis using a HighCapacity cDNA Reverse Transcriptase Kit (Applied Biosystems, Waltham, MA, USA). The cDNA was diluted 16-fold and 5 μl was subjected to real-time quantitative PCR using gene-specific primers (listed in Supplemental Table S9) and SYBR Green PCR master mix in an ABI-7000 sequence detection system (Applied Biosystems). The samples were measured by three repeats.

### β-Galactosidase assays

To test the activity of TFs, we performed one-hybrid or promoter activity assays as described previously (Hsu et al. 2011). The activity of *lacZ*-encoded β-galactosidase was used to represent TF or promoter activity. For the β-galactosidase assay, log-phase cells cultured in the presence of DMSO or 50 μM GdA were harvested and re-suspended in 400 μl Z buffer (0.06 M Na_2_HPO_4_, 0.04 M NaH_2_PO_4_, 0.01 M KCl, 1 mM Mg_2_SO_4_ ・ 7H_2_O, 2.5% β-mercaptoethanol). Next, the cells in 100 μl suspension buffer were lysed by adding 15 μl 0.1% SDS and 30 μl chloroform and then vortexed for 15 seconds. We added 0.2 ml of 4 mg/ml ONPG (o-nitrophenyl-β-galactopyranoside) (N1127, Sigma-Aldrich) substrate into the suspension and vortexed for a further 15 seconds. Since β-galactosidase is activated at 37 ℃, we incubated the suspension at 37 ℃ until the color of the suspension changed to yellow. The reaction was stopped by adding 400 μl 1 M Na_2_CO_3_. Then, we took the supernatant to measure absorbance at 420 and 550 nm. Miller units were calculated to represent β-galactosidase activity, according to the formula 1000*[OD_420_-(1.75*OD_550_)] / [time (min) *volume(ml)*OD_600_]. For each sample, we performed three technical repeats.

## Data access

The datasets generated and analyzed in this study are available in NCBI under the accession number BioProject PRJNA606397.

## Acknowledgements

We thank members of the Leu lab for helpful discussion and comments on the manuscript. We also thank John O’Brien for manuscript editing, and the IMB Genomics core and Academia Sinica Sequencing core for sequencing services. This work was supported by Academia Sinica of Taiwan (grant no. AS-IA-105-L01 to JYL; AS-TP-107-ML06 to JYL and HKT) and the Taiwan Ministry of Science and Technology (MOST 109-2326-B-001-015 to JYL; MOST108-2221-E-001-014-MY3 to HKT).

## Author contributions

JYL conceived the study. PHH, HKT and JYL designed analyses and interpreted results. PHH, CWL and FHK performed the experiments. PHH and HKT performed RNA-seq and computational analysis. PHH and JYL wrote the paper. All authors read and approved the final manuscript.

